# Viral vector-mediated *SLC9A6* gene replacement reduces cerebellar dysfunction in the *shaker* rat model of Christianson syndrome

**DOI:** 10.1101/2024.10.31.621435

**Authors:** Collin J Anderson, Karla P Figueroa, Sharan Paul, Mandi Gandelman, Warunee Dansithong, Joseph A Katakowski, Daniel R. Scoles, Stefan M Pulst

## Abstract

**Background:** Christianson syndrome (CS) is an x-linked recessive neurodevelopmental and neurodegenerative condition characterized by severe intellectual disability, cerebellar degeneration, ataxia, and epilepsy. Mutations to the *SLC9A6* gene encoding NHE6 are responsible for CS, and we recently demonstrated that a mutation to the rat *Slc9a6* gene causes a similar phenotype in the spontaneous *shaker* rat model, which exhibits cerebellar degeneration with motor dysfunction. In previous work, we used the PhP.eB-L7-Slc9a6-GFP adeno-associated viral (AAV) vector to demonstrate that gene replacement in Purkinje cells reduced the *shaker* motor and molecular phenotype.

**Methods:** We carried out a 20-week longitudinal study evaluating the impact of Purkinje cell-specific gene replacement on ataxia and tremor. Taking advantage of the high homology between human *SLC9A6* and rat *Slc9a6*, we tested a more clinically relevant construct, AAV9-CAG-hSLC9A6 AAV vector in the *shaker* rat. In both experimental cohorts, we performed molecular studies to evaluate expression of NHE6 and key cerebellar markers. We then characterized the relationship between molecular markers and motor function, as well between tremor and ataxia.

**Results:** Administration of either of PhP.eB-L7-Slc9a6-GFP or AAV9-CAG-hSLC9A6 AAV vectors led to significant improvement in the molecular and motor phenotypes. The abundance of each disease-relevant cerebellar proteins was significantly correlated to motor ataxia. Further, we found that the relationship between cerebellar ataxia and tremor devolved over time, with disease modifying therapy disrupting their temporal relationship.

**Conclusions:** These findings impact future *SLC9A6*-targeted gene therapy efforts for CS and strongly support gene replacement as a viable therapeutic strategy. Furthermore, tremor and ataxia phenotypes may arise from dissociable cerebellar mechanisms.

**Graphical Abstract:** 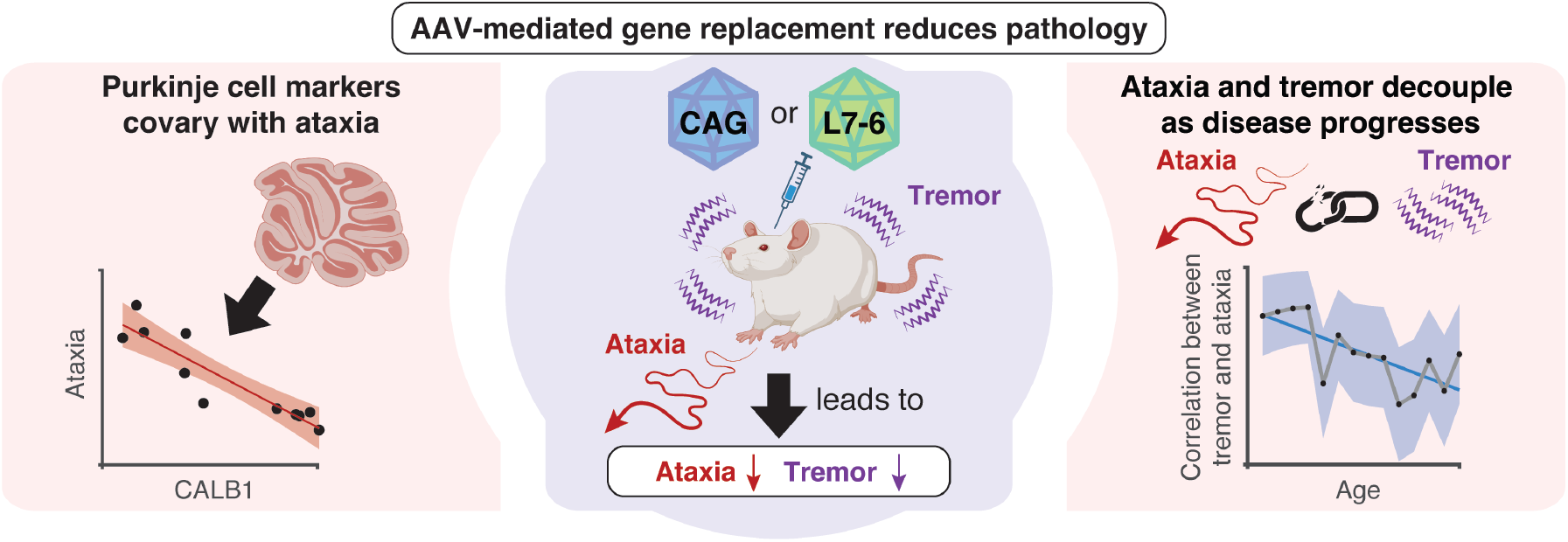

## 1. Background

Christianson syndrome (CS) is an X-linked recessive disorder that presents with developmental delays, intellectual disability, cerebellar degeneration, progressive ataxia, and epilepsy, among other symptoms^1^. CS is caused by loss of function in *SLC9A6* encoding NHE6, a sodium-hydrogen exchanger that regulates pH in early and recycling endosomes^2^. NHE6 is particularly highly expressed in several brain regions and neuronal types, including cerebellar Purkinje cells, coinciding with extensive Purkinje cell degeneration outside of lobule x^3^. CS is untreatable, necessitating therapeutic development.

We previously published on the Wistar Furth *shaker* rat, a spontaneous model of cerebellar degeneration, tremor, and ataxia^4–7^ and demonstrated that this phenotype was caused by a frame-shifting, early-truncating, loss-of-function mutation in *Slc9a6*. This established the *shaker* rat as a spontaneous model of the X-linked condition CS^8^. The *shaker* rat arose spontaneously from a breeding colony at St Louis University in the 1990s ^9–12^, and we have since used the model for gene discovery purposes^4,8^, as well as the development of novel deep brain stimulation approaches aimed at reducing both ataxia and tremor^5^.

In establishing *Slc9a6* as the gene mutated in the *shaker* rat, we performed functional complementation studies that employed adeno-associated viruses (AAVs) to demonstrate the causality of the identified mutation^8^. Previous work focused primarily on gene discovery and replacement and centered on ataxia, which is a primary motor symptom in CS. In this report, we expand on the motor outcomes described in our previous work using PhP-eB-L7-Slc9a6-GFP AAV targeted gene replacement specifically to Purkinje cells, with emphases on the progression of motor symptoms and the complex relationship between tremor and ataxia. Clinical evaluations of tremor and ataxia have traditionally occurred in the context of natural disease progression or with symptomatic relief in the case of tremor, simply based on the lack of disease modifying treatments for either symptom or any form of treatment for ataxia^13,14^. Thus, the preclinical modification of disease progression provides a rare opportunity to study the complex relationship between ataxia and tremor both with and without intervention.

Here we show that AAV-mediated gene replacement in Purkinje cells has significant efficacy in reducing both tremor and ataxia in the *shaker* rat, further supporting gene replacement as a therapeutic strategy for CS. We found that *SLC9A6* replacement was disease modifying, for both tremor and ataxia, but in differential manner, modifying the relationship between tremor and ataxia. Thus, tremor and ataxia in the *shaker* rat may arise from distinct cerebellar mechanisms. Finally, we administered a translationally relevant viral construct, AAV9-CAG-hSLC9A6 not encoding GFP, to the *shaker* rat and again found molecular and motor benefit with broader targeting.

## 2. Methods

All animal work was performed under an approved Institutional Animal Care and Use Committee protocol at the University of Utah, and a total of 50 animals are reported on herein. Rats were housed 2-5 per cage under a standard 12:12 h light cycle with free access to food and water. We maintain the *shaker* colony in the Wistar Furth inbred background (Envigo RMS LLC).

### Genotyping methods

All rats were genotyped as previously described^8^. Briefly, we sought to identify all possible genotype outcomes via PCR using a forward primer of 5′-AAGACATGGCTGTGGCTCGG-3′ and a reverse primer of 5′-AGCTAGGGGACAGGGGTCCG-3′, generating a 363 base pair amplicon. The indel was confirmed via Sanger sequencing utilizing the same above-mentioned forward primer.

### AAV production

PhP.eB-L7-Slc9a6-GFP and PhP.eB-L7-GFP AAV vectors were produced as previous described^8^. Briefly, we PCR-amplified coding sequences from a cDNA library made from wild type rat cerebellar RNA with the forward primer of 5′-TTTATGGCTGTG GCTCGGCGCGGCTGG-3′ and a reverse primer of 5′-TTTCGGCTGGACTGTGTCTTGTGTCATC-3’. We cloned the PCR product into a TOPO vector (Thermofisher, Cat# K457502). We re-amplified the rat *Slc9a6* cDNA from the TOPO vector and cloned into pAAV-L7-6-GFP-WPRE^15^ (Addgene, Plasmid #126462, gifted from Hirokazu Hirai) to generate pAAV-L7-6-Slc9a6-GFP. We generated recombinant AAVs in the University of Utah Drug Discovery Core Facility by co-transfecting HEK-293T cells with either pAAV/L7-6-Slc9a6-GFP or the control pAAV-L7-6-GFP, in addition to pHelper (Stratagene, La Jolla, CA, USA) and pUCmini-iCAP-PHP.eB^16^ (Addgene, plasmid #103005, gifted from Viviana Gradinaru), followed by viral particles purification, concentration, and genomic titer determination.

AAV9-hSLC9A6 AAVs were produced as follows: human cDNA sequence for SLC9A6 (hSLC9A6) (Accession: NM_001042537; Version: NM_001042537.2) corresponding to the NHE6-v1 splice variant (See Supplemental Figure 1 regarding the selection of 701 aa NHE6-v1 vs. 669 aa NHE6-v0) was derived from the NCBI DNA database and used to design primers to PCR-amplify the coding sequences from a cDNA library. The primers set was as follows: SLC9A6-F (*BamHI*): 5’-AATTGGATCCGCCACCATGGCTC GGCGCGGCTGGCGGC-3’ and SLC9A6-R (*EcoRI*): TATCGAATTCTTAGGCTGGA CCATGTCTCGT. To generate hSLC9A6 AAV expression plasmid, pAAV-CAG-GFP plasmid (Addgene plasmid #37825, gifted from Edward Boyden) was modified by deleting GFP with *BamHI* and *EcoRI* restriction enzymes. The amplified hSLC9A6 PCR product was directly cloned into GFP pre-deleted pAAV-CAG-GFP plasmid at *BamHI* and *EcoRI* sites, designated as pAAV-CAG-hSLC9A6. The construct was verified by sequencing. Recombinant AAV particles [pAAV-CAG-GFP (control) and pAAV-CAG-hSLCA6] were generated by Signagen Laboratories, MD, USA (https://signagen.com/).

### AAV administration

We administered 18 rats 2.26 × 10^11^ vg of the control PhP.eB-L7-GFP AAV or PhP.eB-L7-Slc9a6-GFP AAV at 33-37 days of age (average 35 days): three male wild type rats, two female heterozygotes, and thirteen *shaker* rats, specifically six males and seven females hemizygous and homozygous, respectively, for the *Slc9a6* mutation. Two male wild type and two female heterozygotes were kept as uninjected controls. Rats were grouped randomly and AAV administration surgeries were performed as previously described^8^. Briefly, we provided analgesia using 0.1 mL bupivacaine local to the incision and 0.1 mg/kg carprofen subcutaneously daily for 3 days. We anesthetized and maintained rats on 2% isoflurane. We opened to the scalp and dried and performed craniotomies targeting the right lateral ventricle. We waited three min post craniotomy and injected 10 µL of AAV at a rate not exceeding 2 µL / min. After five minutes wait post injection, we withdrew the needle at or slower than 1.0 mm / min. Rats were given a 5-day minimum recovery period.

Further, we administered 23 additional rats – 16 hemizygous males and 8 homozygous females – with AAV9-CAG-hSLC9A6 AAV at 28-41 days of age (average 34 days) by the same procedures with a co-injection of AAV9-CAG-GFP. While ages at injection were more variable in the CAG experiment, average age at injection across groups was insignificantly different by one-way ANOVA (f(5,23)=1.29, p=0.31, η^2^=0.264). In comparison to L7 AAVs administered at only one dose, we administered AAV9-CAG-hSLC9A6 AAV to 6 groups of 4 rats at the following titers: 8.0 × 10^9^, 1.7 × 10^10^, 3.6 × 10^10^, 8.0 × 10^10^, 1.7 × 10^11^, 3.0 × 10^11^ vg. Regardless of the AAV9-CAG-hSLC9A6 AAV dose, all rats were administered 1.74 × 10^10 vg AAV9-CAG-GFP. Notably, one rat from the 1.7 × 10^11^ vg group died 7 weeks post injection, prior to any analyses; therefore, this group is reported with 3 rats.

Note that an L7-6 icosahedral icon is used on all figures referencing experiments carried out with PhP.eB-L7-Slc9a6-GFP AAV, while a CAG icosahedral icon is used on all figures referencing experiments carried out with AAV9-CAG-hSLC9A6 AAV. When both are included, the appropriate icon is applied to relevant sub-figures.

### Motor analyses

Tremor and ataxia were computed as previously described^5–8^. Animals in the L7 experiment were recorded from for 30 minutes weekly from 6 to 25 weeks of age, while animals in the CAG experiment and additional uninjected *shaker* rats were recorded from for 30 minutes at one time point: 14 weeks of age. Rats were placed in the force plate actometer^17–19^ chamber for 30 minutes without any cue or task. The original force plate actometer was rewired through a National Instruments Card and spontaneous ambulation was recorded through center of mass tracking at 1000 Hz using National Instruments LabVIEW.

We imported center of mass data into MATLAB and analyzed straightness of gait during auto-identified rapid movements as a measure of gait ataxia by computing a ratio of distance travelled to displacement. While the minimal value achievable on this ratio is 1.0, minor sway is always present in even healthy animals during walking, and some rapid movements involve rounding of corners. Wild type rats average ∼1.6 on this metric with a tight standard deviation – indeed more than 98.5% of wild type recordings at more than 10 weeks of age yielded averages between 1.4 and 1.8. On the other hand, severely ataxic rats average more than 2.8^5,8^, providing substantial ability to quantify improvement in ataxia.

Tremor was quantified through basic Fourier analyses. We computed Fourier transforms of position over 5-second intervals, then averaged over the recording. Previously^5^, we had averaged Fourier transforms of all age-matched wild type rats and computed tremor for the given rat relative to age-matched wild type rats specifically. However, given that tremor does not significantly change over time in wild type averages^5^, we now more simply integrate the area above a generalized wild type curve averaged across all ages. Tremor strength peaks at ∼5 Hz, with typical increases in power from 3 to 8 Hz. Therefore, we conservatively quantify tremor based on a 2-10 Hz band. Finally, we normalize on a scale in which 0.0 is approximately the maximum tremor score ever present for a wild type rat and 1.0 corresponds to severe tremor. Notably, we have previously determined that heterozygous females do not exhibit quantifiable tremor and do not make measurably less coordinated movements than wild type females. It is not possible to breed both homozygous and wild type females in the same litter; further, given an interest in litter matching across groups, all wild type animals were male, but what was labelled as “wild type” was a combination of wild type male and heterozygous female.

### Tissue collection and protein analyses

Tissue collection took place post motor data collection. In 11 of the 22 rats in the L7 cohort, we retained the left half of the cerebellum for western blot analyses, split evenly across groups, whereas half of the cerebellum was preserved from all animals in the CAG motor experiment. For CAG analyses, we also included an additional 2 age-matched uninjected *shaker* rats and 4 age-matched uninjected wild type rats that did not have motor recordings made. Tissue was flash frozen and homogenized for protein extraction. Protein western blots were run as previously described^8,20^, with protein quantification performed relative to beta-actin (ACTB) or glyceraldehyde-3-phostphate dehydrogenase (GAPDH).

The following primary antibodies were used: NHE6 [(1:5000), Abcam, Cat# ab137185], GAPDH [(1:5000), Cell Signaling, rabbit mAb #2118], ACTB [(1:30,000), Sigma-Aldrich mouse mAb #A3854)], CALB1 [(1:5000), Sigma-Aldrich, mouse mAb #C9848], RGS8 [(1:5000), Novus Biologicals, rabbit pAb #NBP2-20153], and PCP2 [(1:3000), Santa Cruz, mouse mAb #sc-137064]. Secondary antibodies used were HRP-Goat Anti-Rabbit IgG (H+L) [(1:5000), Jackson ImmunoResearch Laboratories, Cat# 111-035-144] and HRP-Horse Anti-Mouse IgG (H+L) [(1:5000), Vector Laboratories, Cat# PI-2000].

### Statistical analyses

Statistical comparisons were made via several methods, the associated test reported with each p-value. First, we used two-sample Student’s t-tests, where n is the number of animals. T-tests were performed one-tailed when previous published data directly supported a hypothesis and two-tailed otherwise. Two-tailed tests are noted as such, and other t-tests should be assumed to be one-tailed. Second, we performed repeated measures ANOVA tests with appropriate post-hoc testing. Finally, we performed Pearson correlation testing to quantify p-values from the correlation coefficient *r*. Notably, for correlation testing, significant correlations are labeled in blue, and insignificant correlations are labeled in red. Further, when correlations are made from individual animals, data points are left unconnected; however, when a data point is generated from a group of animals in correlation analyses across time points, data points are connected.

## 3. Results

All work was completed in Wistar Furth rats, utilizing the Wistar Furth *shaker* rat model of Christianson syndrome with wildtype Wistar Furth rats as controls. Two separate AAV experimental blocks were carried out. First, we administered PhP.eB-L7-Slc9a6-GFP AAV prior to Purkinje cell degeneration and performed longitudinal motor analysis prior to molecular analyses. Second, we administered AAV9-CAG-SLC9A6 at a similar timepoint and quantified motor performance at a single timepoint prior to molecular analyses. The results are combined into three sub-sections. The first pertains to the effects of AAV-mediated gene replacement in the context of CS-relevant motor and molecular dysfunction. The second pertains to the relationship between molecular markers in the cerebellum and motor function. Finally, the third pertains to the relationship between tremor and ataxia.

### 3.1 Gene replacement reduces Christianson syndrome-relevant motor and molecular dysfunction, whether targeted to Purkinje cells or ubiquitously

#### AAV gene replacement targeted to Purkinje cells reduces gait ataxia

In rats administered PhP.eB-L7-Slc9a6-GFP AAV, control PhP.eB-L7-GFP AAV, or uninjected controls at an average of 35 days of age, we tracked ataxia of gait weekly from 6 to 25 weeks of age. We measured gait ataxia by comparing total distance travelled to net displacement in automatically identified, uncued rapid movements, as in previous work^5,7,8^. We performed an ANOVA accounting for time as a variable and found a difference across groups in the context of ataxia, comparing from 10 weeks on, when there was first an ataxia phenotype in untreated *shaker* rats (repeated measures ANOVA, f=184.3, p=1.29 × 10^−66^). In a post-hoc test comparing just *shaker* rats administered PhP.eB-L7-Slc9a6-GFP AAV to those administered PhP.eB-L7-GFP AAV, we found a significant reduction in ataxia in *shaker* rats administered the therapeutic AAV (Repeated measures ANOVA, f=69.7, p=2.04 × 10^−14^) (Figure 1A). The earliest motor symptom onset occurred at 10 weeks of age. We compared the groups of *shaker* rats at individual time points and found that PhpP.eB-L7-Slc9a6-GFP AAV reduced gait ataxia significantly at all time points after 9 weeks of age (Figure 1A) compared to PhP.eB-L7-GFP AAV-injected controls, except for 19 weeks of age (Tukey’s HSD, p<0.05 for all except p=0.0622 at 19 weeks). Further, we compared the total ataxia burden exhibited by each rat from 10 weeks of age on, finding that this was significantly reduced in *shaker* rats administered the PhP.eB-L7-Slc9a6-GFP AAV compared to those administered the control PhP.eB-L7-GFP AAV (student’s t-test, p=0.004806) (Figure 1B).

**Figure 1:**
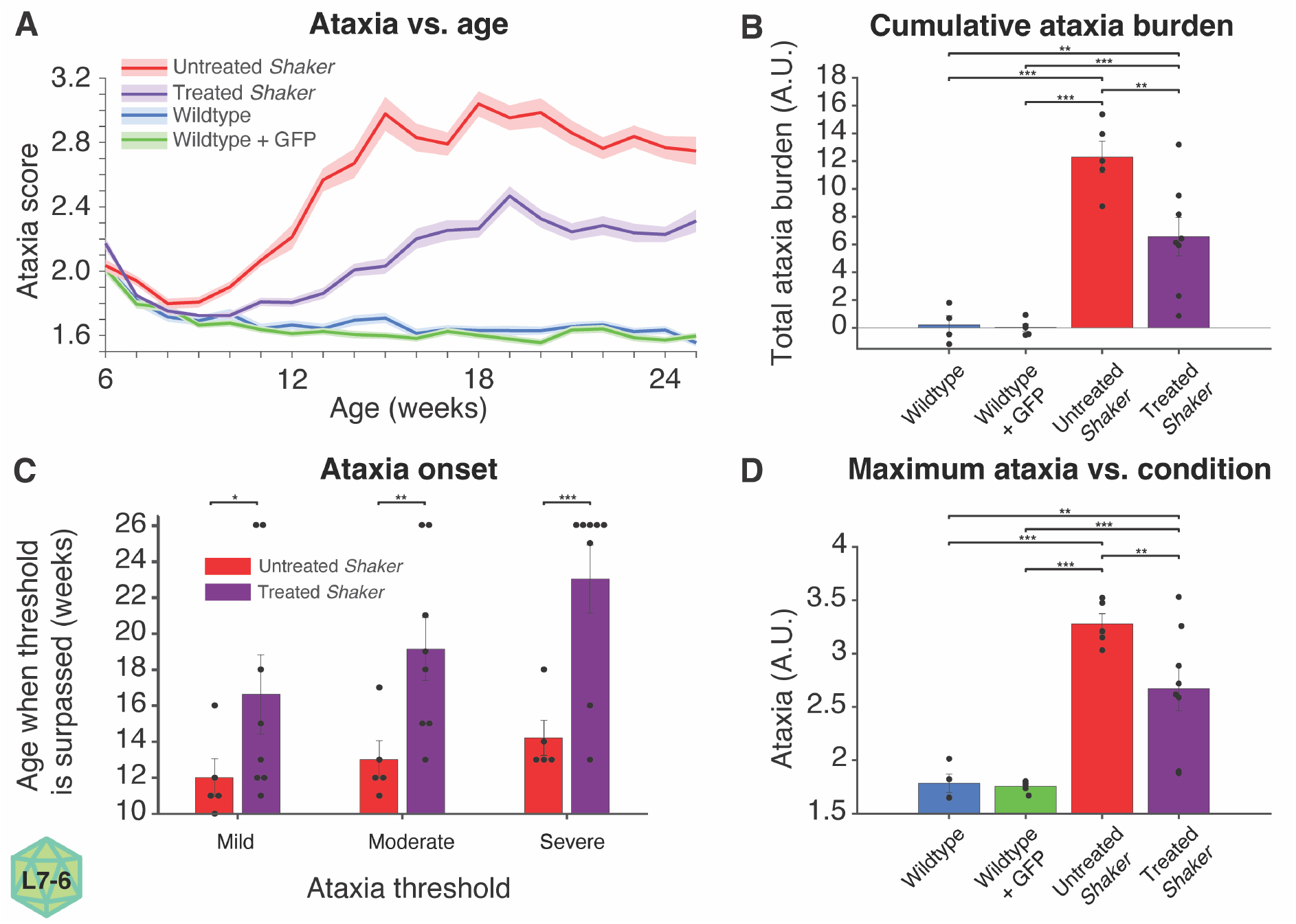
PhP.eB-L7-Slc9a6-GFP AAV reduces ataxia. **A**. Untreated *shaker* rats administered a control PhP.eB-L7-GFP AAV progressively become more ataxic, and PhP.eB-L7-Slc9a6-GFP AAV administered at 5 weeks of age significantly reduced gait ataxia. Administration of a control PhP.eB-L7-GFP AAV to wild type rats had no impact on coordination of gait. **B**. PhP.eB-L7-Slc9a6-GFP AAV decreased cumulative ataxia burden in *shaker* rats by 49.6%, computed as the sum of ataxia scores above 1.6, as calculated in A. **C**. PhP.eB-L7-Slc9a6-GFP AAV delays onset of ataxia in the context of crossing thresholds for mild, moderate, and severe ataxia. **D**. PhP.eB-L7-Slc9a6-GFP AAV decreased the maximal ataxia score experienced by *shaker* rats, as calculated in A.

We compared the onset time for ataxia across groups, given several thresholds. Using the metric described above, wild type rats had average ratios of approximately 1.6, while a threshold of 2.0 represents visually perceivable gait ataxia, 2.4 represents moderate ataxia, and 2.8+ represents severe ataxia. Notably, given that some rats never crossed some or all the ataxia thresholds, we compared the final time points at which an animal hadn’t yet crossed the threshold level of ataxia as a proxy for onset, with a maximum value of 26 weeks, given a final time point of 25 weeks. With ataxia thresholds of each of 2.0, 2.4, and 2.8, we found that PhP.eB-L7-Slc9a6-GFP AAV delayed the onset of ataxia in *shaker* rats (student’s t-test, p=0.0268, p=0.00623, p=0.000979, respectively) (Figure 1C). Finally, we evaluated the peak ataxia exhibited by each rat across all time points, finding that PhP.eB-L7-Slc9a6-GFP AAV decreased peak ataxia significantly (student’s t-test, p=0.0146) (Figure 1D). For all measures in Figure 1, both untreated and treated *shaker* rats were significantly more symptomatic than wild type rats, and no differences were found between uninjected wild type rats and those administered the control AAV.

#### AAV gene replacement targeted to Purkinje cells reduces tremor

We tracked tremor in the above animals from 6-25 weeks of age. We performed a repeated measures ANOVA accounting for time as a variable and found a difference among groups in the context of tremor (repeated measures ANOVA, f=34.1, p=4.45 × 10^-21). In a post-hoc test comparing just *shaker* rats administered PhP.eB-L7-Slc9a6-GFP AAV to those administered PhP.eB-L7-GFP AAV, we found a significant reduction in tremor in *shaker* rats administered the therapeutic AAV (Repeated measures ANOVA, f=13.4, p=0.000354) (Figure 2A). Transduction of Purkinje cells with PhP.eB-L7-Slc9a6-GFP AAV yielded numerally lower tremor at all time points 12 weeks of age and on except for the final time point, 25 weeks of age; however, many time points did not individually yield significant differences via Tukey’s HSD. The total tremor burden exhibited by each rat from 10 weeks of age on was significantly reduced in *shaker* rats administered the PhP.eB-L7-Slc9a6-GFP AAV compared to those administered the control AAV (student’s t-test, p=0.00313) (Figure 2B).

**Figure 2:**
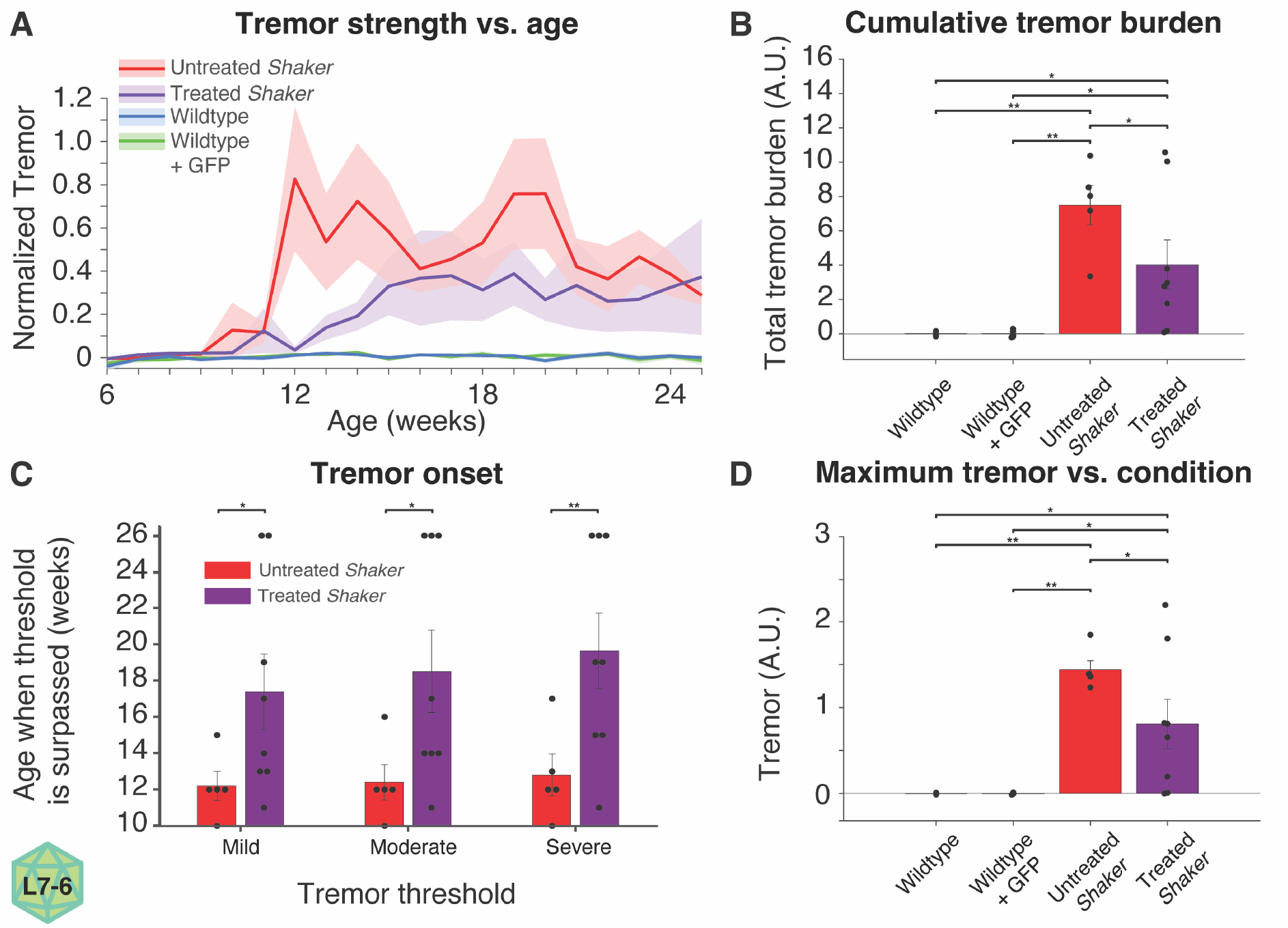
PhP.eB-L7-Slc9a6-GFP AAV reduces tremor. **A**. Untreated *shaker* rats administered a control PhP.eB-L7-GFP AAV display substantial tremor, significantly reduced by the administration of PhP.eB-L7-Slc9a6-GFP at 5 weeks of age. Further, the administration of a control PhP.eB-L7-GFP AAV to wild type rats had no impact in the context of tremor. **B**. PhP.eB-L7-GFP AAV decreased the cumulative tremor burden in *shaker* rats by 39.7%. **C**. PhP.eB-L7-Slc9a6-GFP AAV delayed onset of tremor in the context of crossing thresholds for mild, moderate, and severe tremor. **D**. PhP.eB-L7-Slc9a6-GFP AAV decreased the maximal tremor experienced by *shaker* rats. Notably, as described in the methods, a tremor score of 0.0 represents a typical maximum amount of tremor quantified in wild type rats; therefore, some scores slightly below 0.0 are expected.

We compared the onset time for tremor across groups, given several thresholds. Our normalized numerical scale was designed so that a value of 0 represented no tremor (the maximum 5-8 Hz spectral power a wild type rat would exhibit) while a value of 1 represented the spectral power corresponding to what was qualitatively identified as severe tremor in previous work^5^. Notably, given that some rats never crossed some or all the tremor thresholds, we compared the final time point at which an animal hadn’t yet crossed the threshold level of tremor as a proxy for onset, with a maximum value of 26 weeks, given a final time point of 25 weeks. With a tremor threshold of each of 0.1, 0.25, and 0.5, we found that PhP.eB-L7-Slc9a6-GFP AAV delayed the onset of tremor in *shaker* rats (student’s t-test, p=0.0228, p=0.0174, p=0.00799, respectively) (Figure 2C). Finally, we evaluated the peak tremor exhibited by each rat across all timepoints, finding that within *shaker* rats, PhP.eB-L7-Slc9a6-GFP AAV decreased peak tremor significantly (student’s t-test, p=0.0.0350) (Figure 2D). For all measures in Figure 2, both untreated and treated *shaker* rats were significantly more symptomatic than wild type rats, and no differences were found between uninjected wild type rats and those administered the control AAV.

#### AAV gene replacement with ubiquitous targeting reduces gait ataxia and molecular markers of cerebellar health

Following the above L7 AAV experiments, we began work with a more translationally relevant viral construct. Compared to the PhP-eB.L7-Slc9a6-GFP AAV, the AAV9-CAG-hSLC9A6 AAV has 4 primary differences: removal of the GFP tag, replacement of the rat *Slc9a6* gene with the human *SLC9A6* gene, the use of the ubiquitously expressing CAG promoter instead of the Purkinje cell-specific L7-6 promoter, and the use of the more clinically relevant AAV9 instead of PhP.eB. We administered varying doses of AAV9-CAG-hSLC9A6 AAV – ranging from 0.8 to 30 × 10^10 vg – at an average of 34 days of age and measured ataxia and tremor at 14 weeks of age before immediately collecting cerebellar tissue for molecular analyses. We found that cerebellar NHE6 expression was significantly increased in rats administered AAV9-CAG-hSLC9A6 AAV (student’s t-test, p=2.68 E-9) (Figure 3A-B), as was cerebellar CALB1 expression (student’s t-test, p=0.000347) (Figure 3C-D). AAV9-CAG-hSLC9A6 AAV administration yielded a reduction in ataxia (student’s t-test, p=0.0347) (Figure 3E). However, while AAV-CAG-hSLC9A6 AAV administration trended towards reduced tremor, this was not significant (student’s t-test, p=0.199) (Figure 3F), matching the wide variability in tremor at similar time points in the L7 experiments. Note that molecular outcomes from the above L7 experiments matching those in Figure 3 A-D are not reported as they have previously been published.^8^

**Figure 3:**
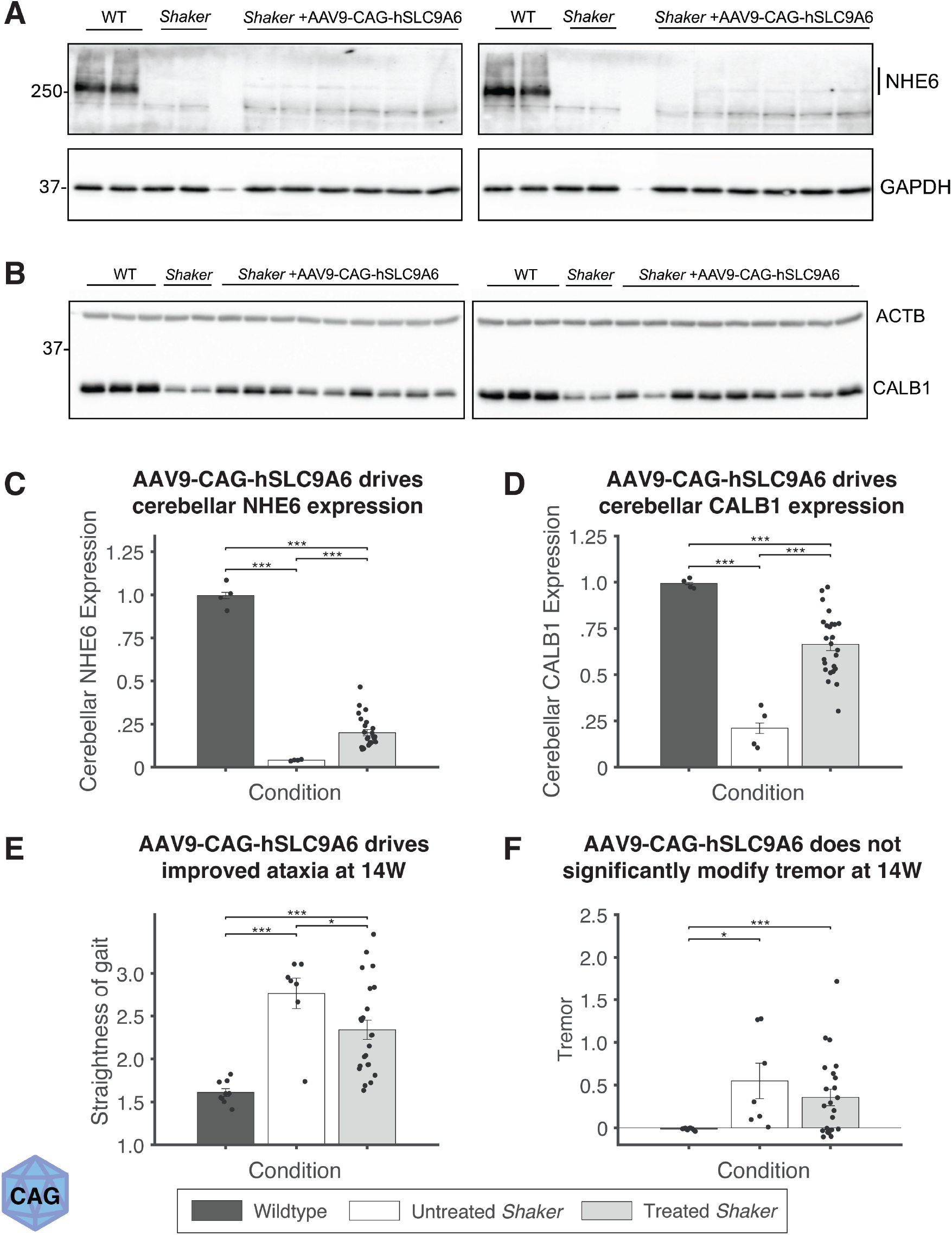
AAV9-CAG-hSLC9A6 AAV increased cerebellar NHE6 and CALB1 abundances determined by western blotting and decreased motor ataxia. **A-D**. Mirroring findings with PhP.eB-L7-Slc9a6-GFP AAV, administration of AAV9-CAG-hSLC9A6 AAV at ∼5 weeks of age minimally increased NHE6 expression (**A, C**), but this minimal increase led to proportionally greater increases in CALB1 expression (**B, D**). **E**. AAV9-CAG-hSLC9A6 AAV administration at 5 weeks of age significantly improved motor ataxia at 14 weeks of age compared to 7 untreated *shaker* rats (5 administered a control AAV and 2 uninjected). **F**. Rats treated with AAV9-CAG-hSLC9A6 trended towards a decrease in tremor at 14 weeks of age, albeit with substantial variability. Note that each data point is an individual rat, and all n-values for statistics are calculated as number of rats, not technical replicates.

Finding evidence of therapeutic benefit with L7- and CAG-based AAVs in this and previous^8^ work, we took advantage of the variable nature of therapeutic benefit to ask fundamental questions: how do molecular markers co-vary with motor dysfunction, and how are ataxia and tremor interrelated in the context of CS-related cerebellar dysfunction?

### 3.2 The abundance of key cerebellar health markers is strongly tied to ataxia, but less so to tremor

#### In the context of AAV9-CAG-hSLC9A6 AAV, NHE6 and CALB1 expression are correlated with ataxia, but not tremor

As shown in Figure 3 above, all *shaker* rats administered AAV9-CAG-hSLC9A6 AAV exhibited stronger NHE6 expression than uninjected *shaker* rats. However, in the context of those treated with AAV9-CAG-hSLC9A6 AAV, viral dose and NHE6 expression were not significantly correlated (Pearson test, r=0.0403, p=0.855) (Supplemental Figure 2A). Similarly, AAV dose was not correlated with CALB1 (Pearson test, r=-0.1668, p=0.4469) (Supplemental Figure 2B), ataxia (Pearson test, r=0.1586, p=0.4698) (Supplemental Figure 2C), or tremor (Pearson test, r=0.0959, p=0.6631) (Supplemental Figure 2D). However, NHE6 was correlated with CALB1 (Pearson test, r=0.4920, p=0.0125) (Figure 4A) and ataxia (Pearson test, r=-0.4211, p=0.0361) (Figure 4B). Further, CALB1 was highly correlated with ataxia (Pearson test, r=-0.6704, p=0.000247) (Figure 4C). Interestingly, tremor was not significantly correlated with either NHE6 (Pearson test, r=-0.3056, p=0.1374) (Figure 4D) or CALB1 (Pearson test, r=-0.3578, p=0.0791) (Figure 4E).

**Figure 4:**
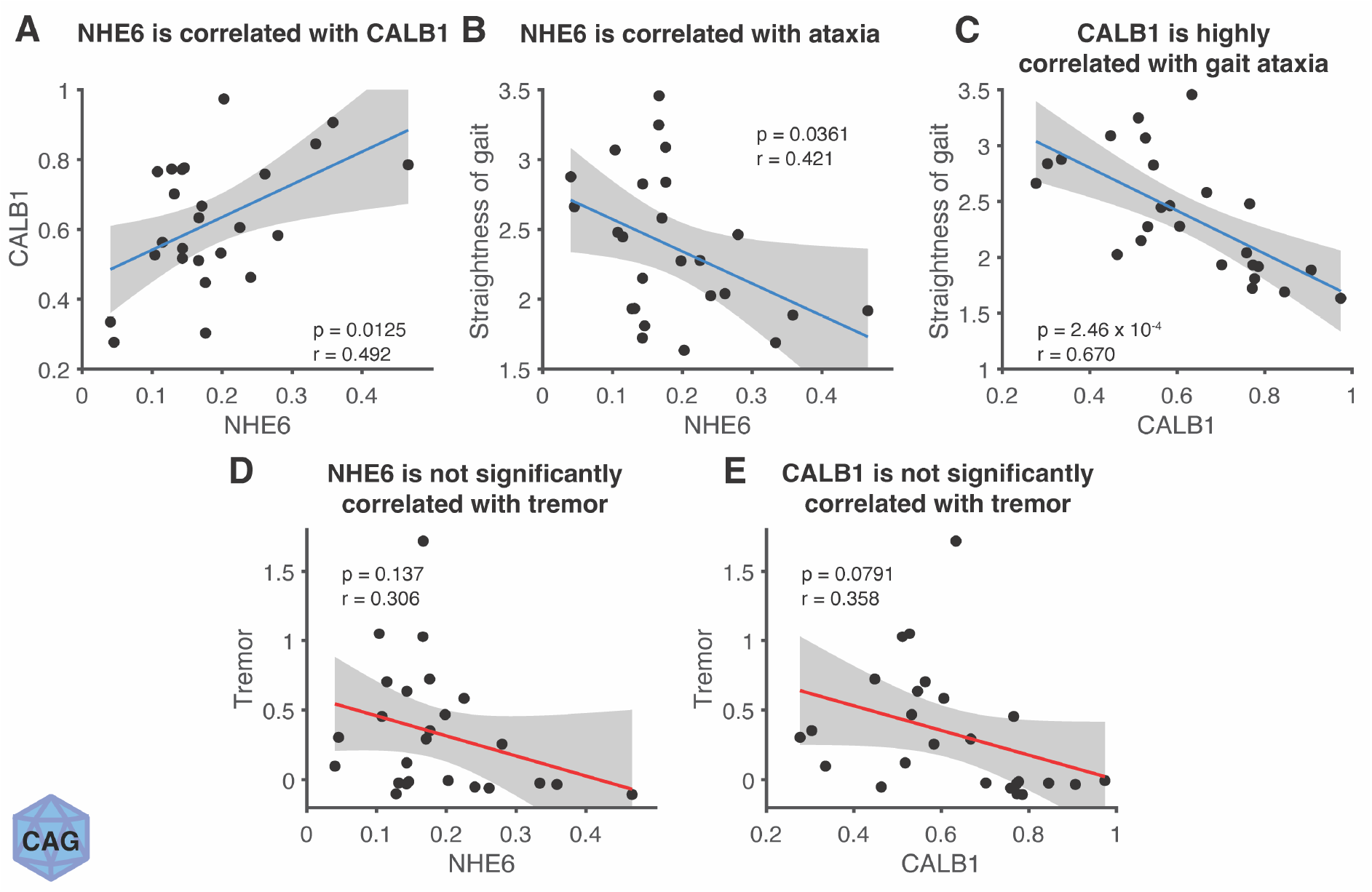
Following AAV9-CAG-hSLC9A6 AAV administration, ataxia, NHE6, and CALB1, are interrelated, but NHE6 and CALB1 are not correlated with tremor at 14 weeks of age. **A**. NHE6 abundance following viral administration of AAV9-CAG-hSLC9A6 AAV positively correlates with CALB1 expression, suggesting a rescue of Purkinje cells. **B**. NHE6 abundance is likewise negatively correlated with gait ataxia. **C**. CALB1 is highly negatively correlated with gait ataxia. **D-E**. While NHE6 and CALB1 abundance were correlated with ataxia and ataxia is correlated with ataxia, NHE6 and CALB1 are not significantly correlated with tremor.

#### In the context of PhP.eB-L7-Slc9a6-GFP AAV, expression of NHE6 and key cerebellar proteins is correlated with gait ataxia, but not with tremor

Following the above analyses in the context of AAV9-CAG-hSLC9A6, we returned to the analysis of PhP.eB-L7-Slc9a6-GFP AAV to evaluate the relationship between motor outcomes and various cerebellar health-relevant expression levels. In a previous report on our PhP.eB-L7-Slc9a6-GFP AAV work, we showed that each of NHE6, CALB1, PCP2, and RGS8 were significantly reduced in *shaker* rats compared to wild type or *Slc9a6* heterozygous rats, with *Slc9a6* significantly reduced from wild type rats to heterozygotes.^8^ Pooling all animals in the L7 AAV study with western blotting data (n=11), we find that each of these metrics is significantly correlated with late-stage coordination, averaged from 21-25 weeks of age (Pearson test, r=-0.732, p=0.0104 for NHE6; r=-0.926, p=0.000043 for CALB1; r=-0.870, p=0.00050 for PCP2; r=-0.928, p=0.000038 for RGS8) (Figure 5A). Notably, these relationships held even when evaluating only within *shaker* rats, except for NHE6, which trended towards significance (Pearson test, r=-0.616, p=0.140).

**Figure 5:**
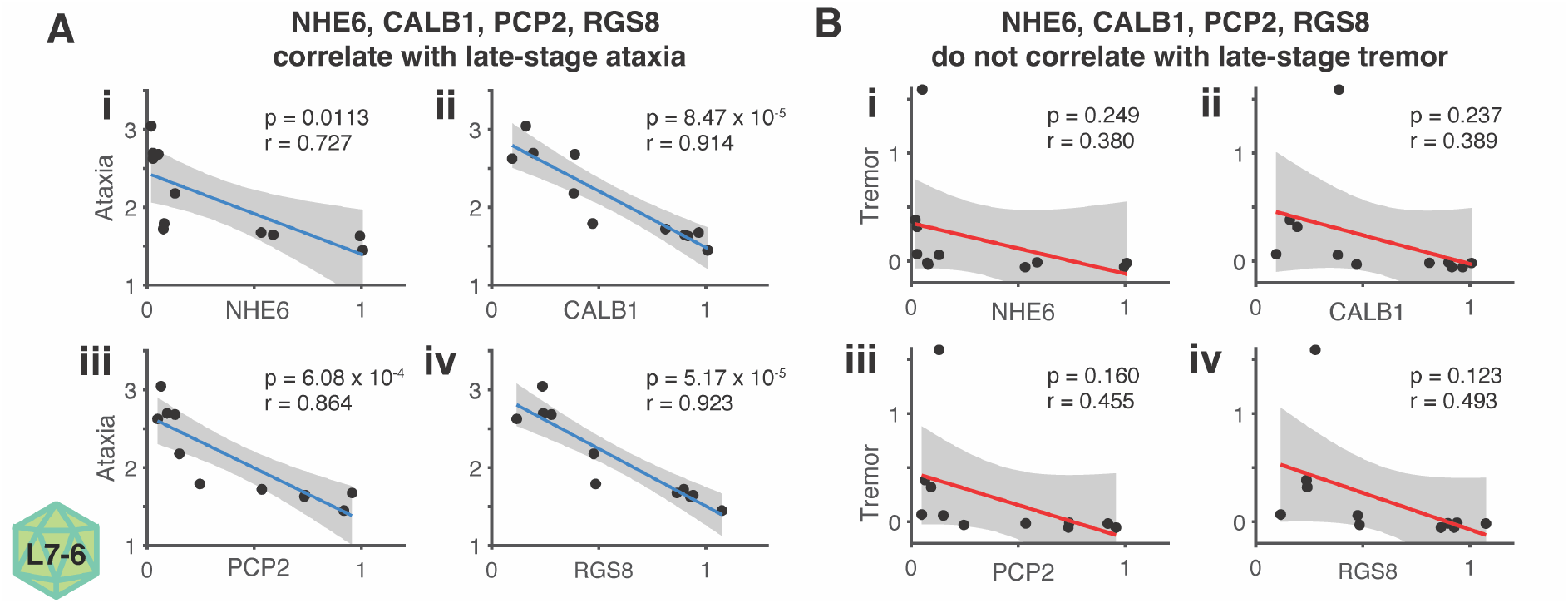
In PhP.eB-L7-Slc9a6-GFP AAV experiments, cerebellar molecular markers are correlated with gait ataxia. **A**. NHE6, CALB1, PCP2, and RGS8 are significantly correlated with late-stage ataxia (21-25 weeks). **B**. Similar correlations are not found with late-stage tremor.

Late-stage tremor, averaged from 21-25 weeks of age, did not significantly correlate with any of these metrics when pooling all animals (Pearson test, -0.493 < r < -0.380, 0.123 < p < 0.249 for all) (Figure 5B). When evaluating only within *shaker* rats, trending relationships weakened further (Pearson test, -0.403 < r < -0.239, 0.371 < p < 0.605).

### 3.3 Tremor and ataxia are interrelated but dissociable

#### Tremor and ataxia are linked, but their relationship weakens over time

In the context of AAV9-CAG-hSLC9A6 AAV experiments, we found that at 14 weeks of age – the single time point of motor recordings – tremor and ataxia were highly correlated (r=0.8474, p=9.06 E-8) (Figure 6A). Thus, we longitudinally analysed the relationship between tremor and ataxia in all *shaker* rats in PhP.eB-L7-Slc9a6-GFP AAV experiments. We analyzed the relationship between tremor and ataxia from 12 weeks on – in other words, after significant symptom onset in both symptoms. In the initial weeks following symptom onset – 12 to 15 weeks – ataxia was highly correlated with tremor (Pearson test, 0.7496<r<0.7491, 0.00119<p<0.00318). However, the correlation between ataxia and tremor decreased over time (Pearson test, r=-0.7274, p=0.00320) (Figure 6B).

**Figure 6:**
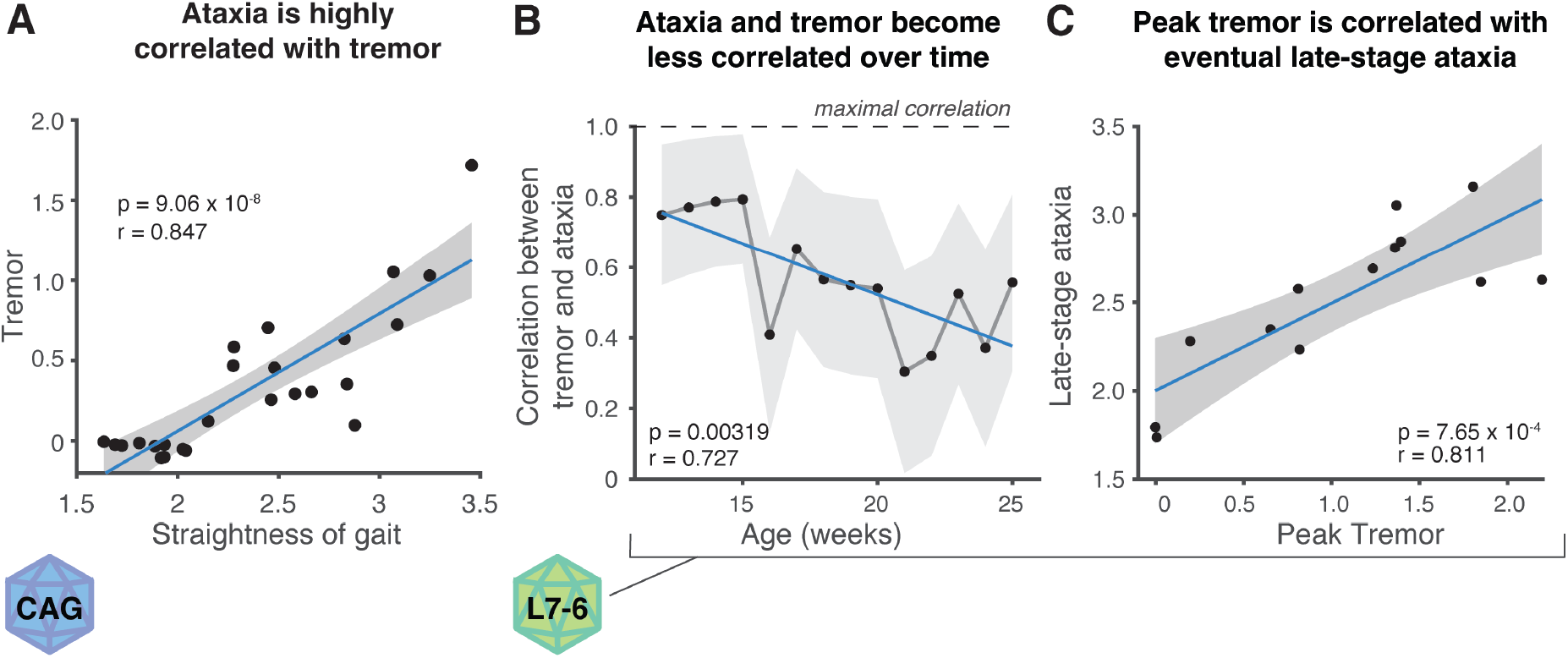
**A**. In the context of AAV9-CAG-hSLC9A6 AAV experiments, ataxia is strongly correlated with tremor at 14 weeks. **B-C**. In the context of PhP.eB-Slc9a6-GFP AAV experiments, the correlation between peak tremor and late-stage ataxia is stronger than the correlation between tremor and ataxia at any given timepoint. **B**. Ataxia and tremor are initially strongly related following symptom onset, but this correlation significantly decreases over time. **C**. Peak tremor occurred prior to 20 weeks in all but 2 animals; however, the correlation between peak ataxia and late-stage ataxia (21-25 weeks) was higher than the correlation between tremor and ataxia at any individual timepoint.

For all but one time point after 18 weeks of age, no significant relationship was found between tremor and ataxia. Peak tremor was reached prior to 20 weeks of age in all but 2 of the 13 *shaker* rats in the study; however, when comparing peak tremor to late-stage ataxia (weeks 21-25), we found that peak tremor was highly correlated with eventual late-stage ataxia (Pearson test, r=0.8112, p=0.000764), achieving a stronger correlation than that between tremor and ataxia at any individual time point (Figure 6C).

#### PhP.eB-L7-Slc9a6-GFP AAV disrupted the temporal relationship between tremor and ataxia

We determined the onset time of tremor and ataxia in all animals based on crossing minimal symptomatic thresholds (0.1 for tremor, 2.0 for ataxia). While onset times for both symptoms were significantly different between treated and untreated shaker rats (Figures 1C, 2C), no difference was found between the onset time for tremor and the onset time for ataxia in either group (2-tail student’s t-test, p=0.883 for untreated, p=0.807 for treated) (Figure 7A). No difference was found between untreated and treated groups in terms the age at which animals reached peak ataxia (defined as at least 90% of their lifetime maximal value) (two-tailed student’s t-test, p=0.8513); however, treated *shaker* rats reached peak tremor values at a later age than untreated *shaker* rats (two-tailed student’s t-test, p=0.0437) (Figure 7B). Peak ataxia occurred significantly later than peak tremor in untreated *shaker* rats (two-tailed student’s t-test, p=0.0402), while no difference was found in treated *shaker* rats (two-tailed student’s t-test, p=0.917). Further, a difference was found across groups, with significantly less delay from peak tremor to peak ataxia in treated *shaker* rats, -0.125 weeks on average, than untreated *shaker* rats, 4.4 weeks on average (two-tailed student’s t-test, p=0.0398) (Figure 7B). Removing animals that didn’t meet a symptomatic threshold (0.1 for tremor, 2.0 for ataxia), significance was no longer achieved, but a strong trend remained (two-tail student’s t-test, p=0.0574).

**Figure 7:**
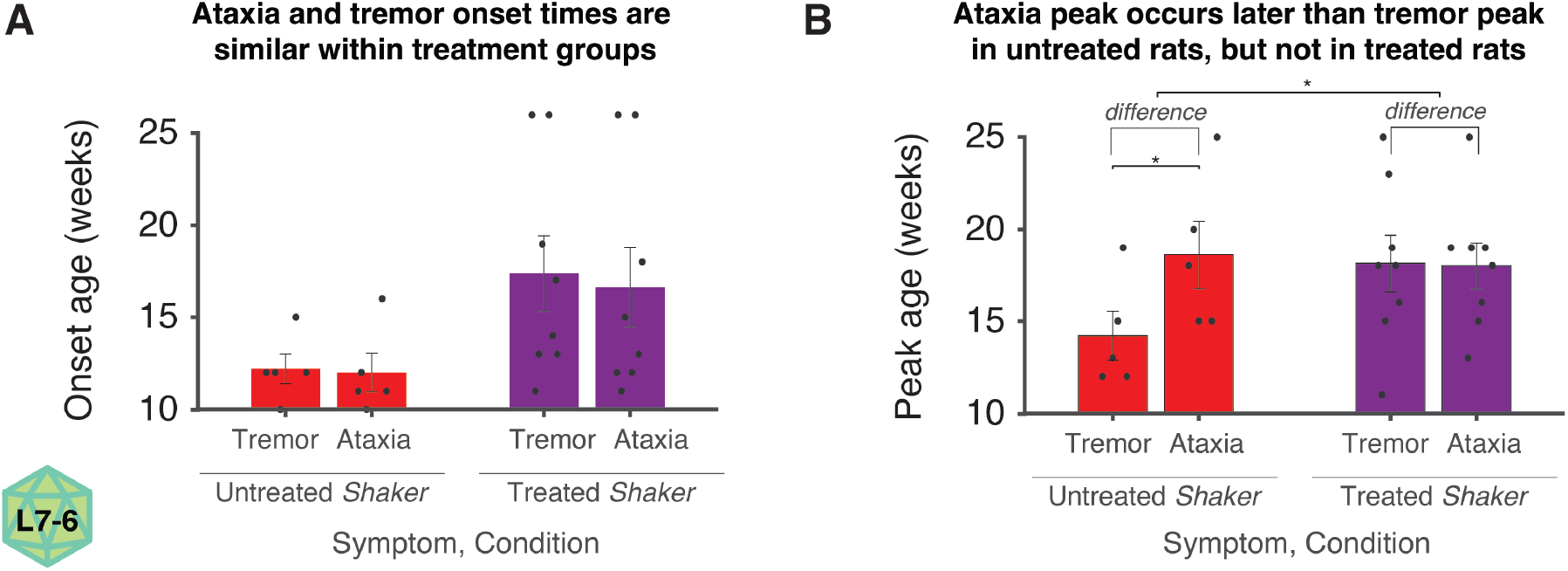
PhP.eB-L7-Slc9a6-GFP AAV disrupts the temporal relationship between tremor and ataxia. **A**. Within treatment groups, tremor and ataxia onset occurs at similar ages. **B**. In untreated *shaker* rats, a significant delay exists from tremor peak to ataxia peak, but this delay is eliminated in treated *shaker* rats.

#### Post-experimental abundance of NHE6 becomes more linked to ataxia over time, but its relationship with tremor does not evolve over time

The degree to which post-experimental NHE6 expression predicted ataxia increased over time (Pearson test, r=-0.7228, p=0.000318) (Figure 8A). In other words, while there is some variability in the timing of ataxia onset in the absence of NHE6, level of NHE6 is highly correlated with the eventual end-stage ataxic burden. Interestingly, such a relationship did not hold for tremor, and a trend was found that post-experimental NHE6 was better at predicting earlier-stage tremor than later stage tremor (Pearson test, r=0.4114, p=0.0715) (Figure 8B).

**Figure 8:**
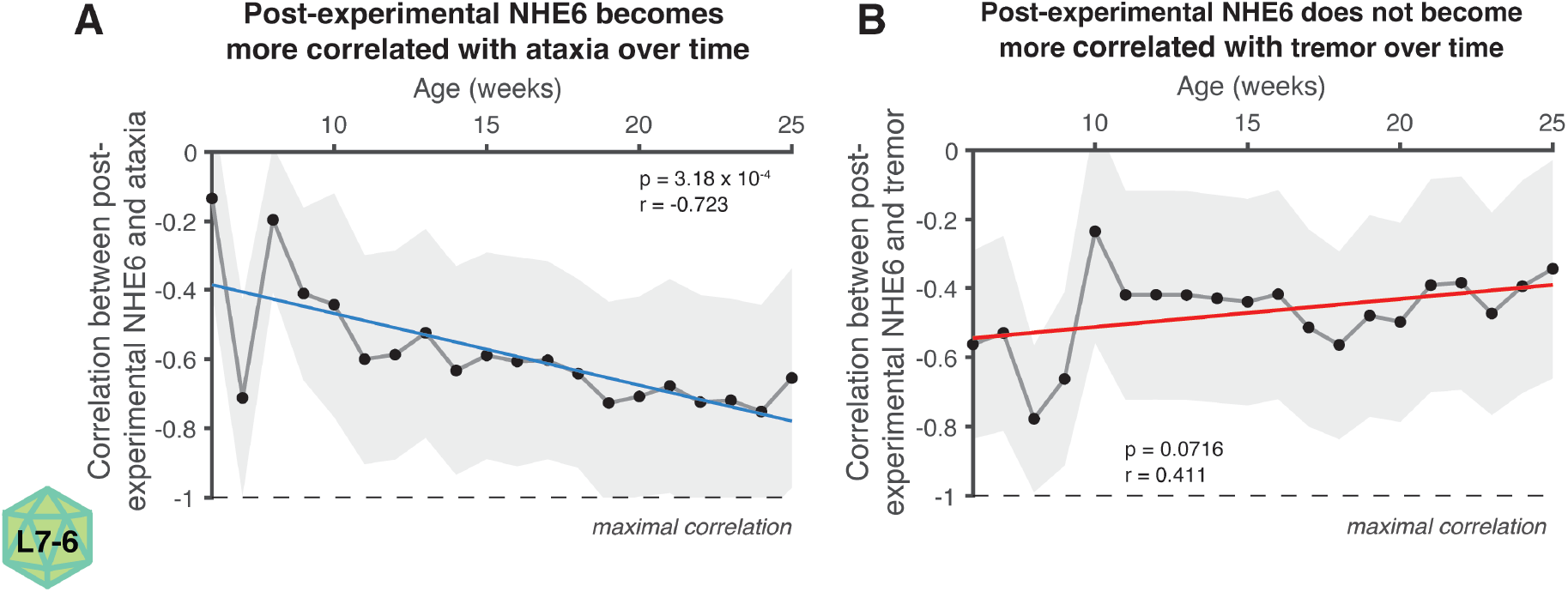
**A**. NHE6 and ataxia were negatively correlated at all timepoints. This negative correlation becomes stronger over time. **B**. NHE6 and tremor were negatively correlated at all timepoints; however, the correlation between NHE6 and tremor does not strengthen over time and trended towards weakening.

## 4. Discussion

Christianson syndrome is characterized by intellectual disability, epilepsy, behavioral abnormalities, and progressive motor dysfunction driven by cerebellar degeneration. Because *SLC9A6* mutations cause loss of NHE6 function, viral vector-mediated gene replacement may be an appropriate therapeutic approach for CS. Here, we demonstrate that multiple viral constructs are capable of decreasing cerebellum-related pathology, and with both constructs, relatively minimal NHE6 expression was required for substantial motor and molecular benefit. These major benefits driven by minimal NHE6 expression match case reports in human patients in which even a minimal degree of functional NHE6 expression is associated with a large decrease in symptomatic burden.^21^ Further, NHE6 loss of function leads to an over-acidification of the endosome^22^, with CS mutations decreasing endosomal pH by 0.1-0.2^23^. Low pH enhances viral release from the endosome^24^, and this potentially may allow CS therapeutics using relatively low viral titers. Ultimately, the combination of our results, human findings, and the potential for decreased titers due to endosomal over-acidification all create reasons for optimism regarding viral gene replacement as a therapeutic strategy for CS. Indeed, treated rats in longitudinal studies had an average overall ∼50% reduction in total ataxia burden and a ∼40% reduction in total tremor burden. These results are promising, but motor dysfunction is only one component of CS symptomology, necessitating substantial additional study.

Beyond CS, these results are informative in a broader discussion of cerebellum-driven motor symptoms. Cerebellar dysfunction leads to numerous movement symptoms, including ataxia and tremor, and the relationship between tremor and ataxia is complex. Tremor is associated with numerous spinocerebellar ataxias^25–27^ and gait/balance issues are associated with multiple forms of tremor^28–30^. In some degenerative ataxias, the presence of specific forms of tremor may be a predictive factor in the rate of ataxia progression, albeit with heterogeneity. For example, SCA1 and SCA6 patients with postural tremor experience slower progression of ataxia, while SCA2 patients with postural tremor experience faster progression of ataxia^25^. In some disorders that regularly feature both tremor and ataxia such as Fragile X-associated Tremor/Ataxia Syndrome (FXTAS), either tremor or ataxia can onset first with a similar likelihood.^31^ It has been proposed that essential tremor may represent an intermediate form of spinocerebellar degeneration^32–35^. However, others have reported that Purkinje cell loss is not a feature of essential tremor^36,37^, generating debate.^38,39^ Increased understanding of the link between tremor and ataxia – as well as that between tremor and cerebellar degeneration – is needed, both for tremor patients and ataxia patients.

Several potential caveats should be addressed. First, and most obviously, we did not find any dose dependence in the context of any molecular or motor outcomes in CAG AAV studies. Notably, this is likely tied to higher doses not associated with stronger NHE6 expression. Thus, we suspect that these findings were related to the differential volumes of injection at different doses, based on previous results showing that increased fluid volume leads to decreased transduction^40^. Indeed, we did not dilute lower doses to match volumes, and it is our hypothesis that doing so in future studies would, in fact, lead to dose-dependent relationships. Second, we note that statistical assumptions were violated in several correlation analyses, with several distributions (NHE6 and tremor in L7 studies) failing a Lilliefors test for normality. While some of these distributions would have met requirements if evaluated on a log scale, a log scale-based evaluation of, say, tremor would not be meaningful; regardless, conclusions would only change minimally if log scales were used for analysis. Finally, it is worth noting that it has been reported that NHE6 expression does not always correspond with areas that exhibit the greatest pathologies. Intriguingly, while NHE6 expression is strong in Purkinje cells, matching Purkinje cell pathology and degeneration, substantial dysfunction is reported in specific regions of hippocampus and amygdala that don’t feature strong endogenous NHE6 expression.^3^ This may complicate gene replacement therapies, necessitating additional study in contexts broader than the cerebellum.

While this reported work is specific to Christianson syndrome, our results find agreement with recent reports that cerebellar tremor and ataxia arise from distinct and dissociable mechanisms^41^. Several points relevant to the question of shared causality of tremor and ataxia merit discussion. First, gene replacement modified the temporal relationship between tremor and ataxia. If tremor were simply based exclusively on PC loss and solely an early motor outcome of Purkinje cell loss that eventually evolves into ataxia, modest rescue of PCs should not have eliminated the substantial separation between tremor peak and ataxia peak present in untreated animals in both this and previous work^4,5^. Second, it is interesting that each molecular marker analyzed was correlated with late-stage ataxia, and most were correlated with peak tremor – which occurred relatively early in symptomatic progression in nearly all cases – but none were significantly correlated with late-stage tremor. Further, we discovered that the degree to which each of the reported molecular markers predicted gait ataxia increased over time, while their relationship with tremor either remained unchanged or weakened over time. Finally, while peak tremor – typically at early time points – was related to eventual end-stage ataxic burden with a significant relationship between tremor and ataxia at early time points, this direct correlation degraded with age. All of these taken together suggest that while the mechanisms underlying the generation of tremor and ataxia in the *shaker* rat are interrelated, they are at most only partially shared and likely dissociable.

Christianson syndrome is a complex neurodevelopmental and neurodegenerative disorder caused by loss of function in the *SLC9A6* gene. In this work, we found strong evidence that viral gene therapy can reduce the ataxic component and related cerebellar dysfunction in CS, but much work remains in the context of other symptoms, dose timing, and further optimization. This work presented the ability to critically evaluate the complex relationship between tremor and ataxia under the rare context of disease-modifying therapy. While the findings may be specific to Christianson syndrome, they may add to the complex discussion of tremor and ataxia.

## Supplemental Figures

**Supplemental Figure 1:**
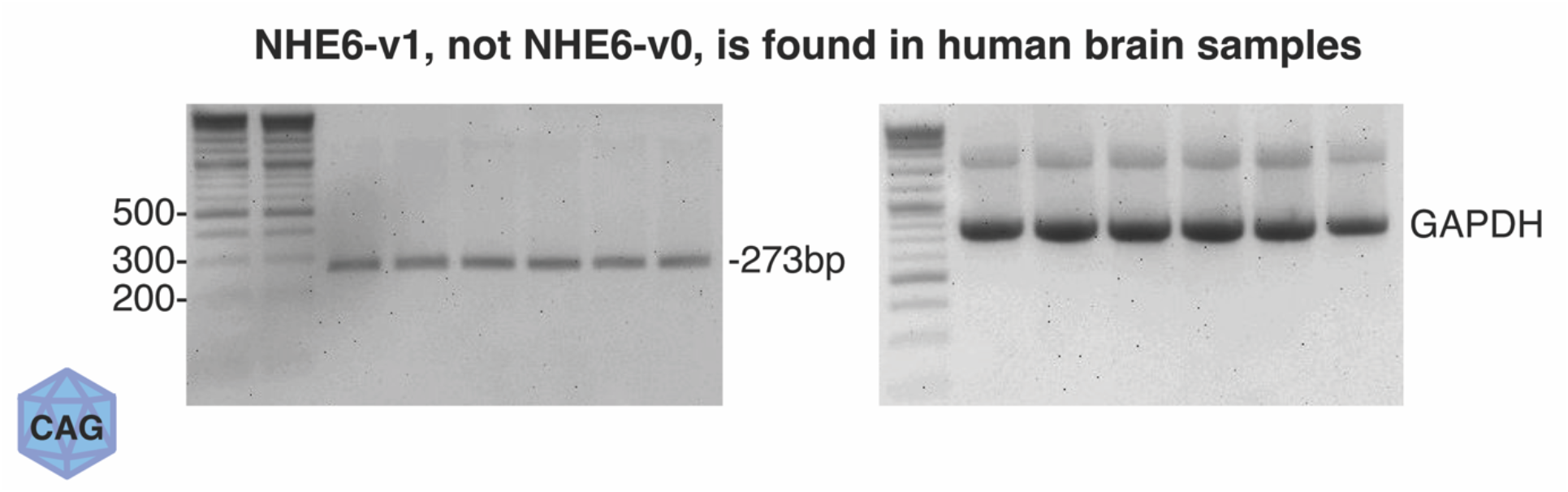
We pooled samples from human brains taken from healthy individuals. We created sets of primers such that NHE6-v0 would generate a 177bp PCR product and NHE6-v1 would generate a 273bp PCR product. We performed RT-PCR and found that the 701 amino acid splice variant (NHE6-v1) is predominant in brain, with the 669 amino acid splice variant (NHE6-v0) not detected. Hence, for AAV9-CAG-hSLC9A6 AAV experiments, we utilized NHE6-v1.

**Supplemental Figure 2:**
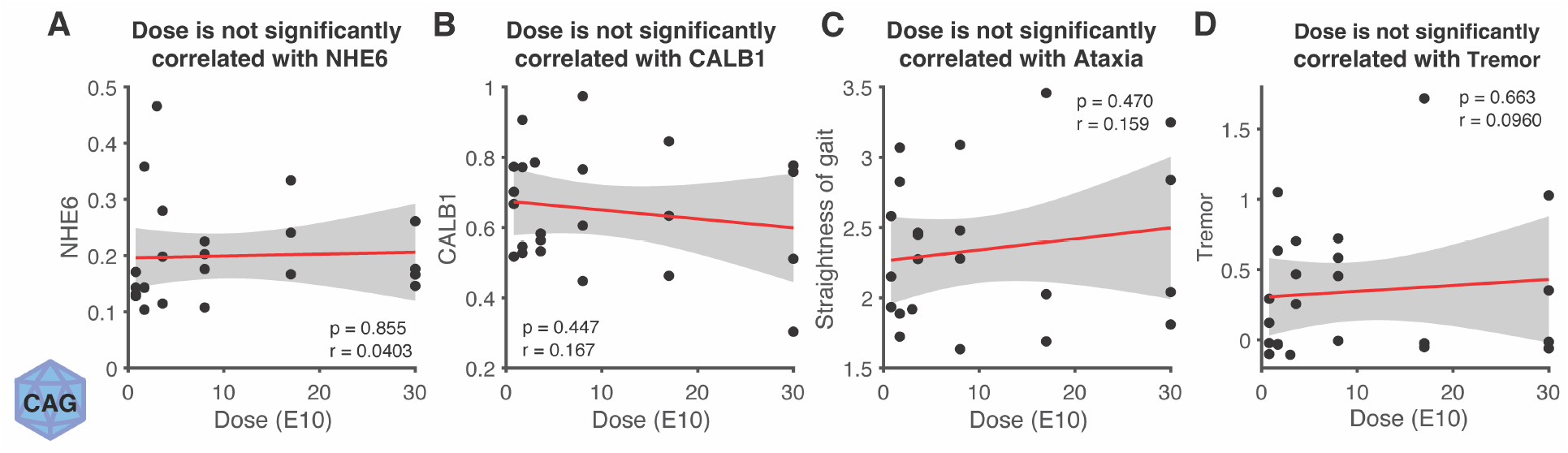
In the context of AAV9-CAG-hSLC9A6, dose (E10) was unrelated to NHE6 (A), CALB1 (B), ataxia (C), or tremor (D). Notably, this AAV9 was delivered at the same concentration regardless of dose, and thus, dose and injection volume covaried. We hypothesize that a lack of dose response but an overall consistent response compared to untreated animals was driven by differences in volume, with less effective transduction with higher volumes.

## Declarations

### Availability of data and materials

All data are available from the corresponding author on reasonable request.

### Funding

NIH NINDS R21 NS104799 (SMP)

NIH NINDS R35 NS127253 (SMP)

3x RTW Charitable Foundation Research Grant (1 to CJA, SP, SMP; 1 to CJA and SMP; 1 to CJA)

University of Sydney Bright Ideas Grant (CJA)

University of Sydney CAPEX Grant (CJA)

## Acknowledgments

The authors thank Daria Nesterovich Anderson for advice on organisation of results and assistance with data visualization.

